# Multiscale Simulation Reveals Passive Proton Transport Through SERCA on the Microsecond Timescale

**DOI:** 10.1101/2020.06.17.157396

**Authors:** Chenghan Li, Zhi Yue, L. Michel Espinoza-Fonseca, Gregory A. Voth

## Abstract

The sarcoplasmic reticulum Ca^2+^-ATPase (SERCA) transports two Ca^2+^ ions from the cytoplasm to the reticulum lumen at the expense of ATP hydrolysis. In addition to transporting Ca^2+^, SERCA facilitates bidirectional proton transport across the sarcoplasmic reticulum to maintain the charge balance of the transport sites and to balance the charge deficit generated by the exchange of Ca^2+^. Previous studies have shown the existence of a transient water-filled pore in SERCA that connects the Ca^2+^-binding sites with the lumen, but the capacity of this pathway to sustain passive proton transport has remained unknown. In this study, we used the multiscale reactive molecular dynamics (MS-RMD) method and free energy sampling to quantify the free energy profile and timescale of the proton transport across this pathway while also explicitly accounting for the dynamically coupled hydration changes of the pore. We find that proton transport from the central binding site to the lumen has a microsecond timescale, revealing a novel passive cytoplasm-to-lumen proton flow beside the well-known inverse proton countertransport occurring in active Ca^2+^ transport. We propose that this proton transport mechanism is operational and serves as a functional conduit for passive proton transport across the sarcoplasmic reticulum.

**SIGNIFICANCE:** Multiscale reactive molecular dynamics combined with free energy sampling was applied to study proton transport through a transient water pore connecting the Ca^2+^-binding site to the lumen in SERCA. This is the first computational study of this large biomolecular system that treats the hydrated excess proton and its transport through water structures and amino acids explicitly. When also correctly accounting for the hydration fluctuations of the pore, it is found that a transiently hydrated channel can transport protons on a microsecond timescale. These results quantitatively support the hypothesis of the proton intake into the sarcoplasm via SERCA, in addition to the well-known proton pumping by SERCA to the cytoplasm along with Ca^2+^ transport.

## INTRODUCTION

The sarcoplasmic reticulum Ca^2+^-ATPase (SERCA) pump is a critical component of Ca^2+^ transport in cells and is also an extensively studied member of the large family of P-Type ATPases. SERCA plays a central role in muscle contraction and intracellular Ca^2+^ homeostasis by clearing cytosolic Ca^2+^. At the cost of one ATP hydrolyzed, SERCA pumps two Ca^2+^ from the cytoplasm into sarcoplasmic reticulum lumen and, at the same time, transports two or three protons in the opposite direction due to the need for charge balance of the binding site in the absence of Ca^2+^ (1-7). During its functional cycle, SERCA prominently populates two types of states, a cytoplasmic facing E1 state and a luminal facing E2 state (8-10). The electroneutrality across the ER membrane during the Ca^2+^ intake is compensated by the proton counter-transport of SERCA as well as other fluxes of ions abundant in the cell, such as K^+^, Na^+^, and Cl^-^ (11). Among them, the Cl^-^ influx was reported to play an essential role in balancing luminal positive charges (12, 13). Several chloride-channel (ClC) family proteins, which were identified as Cl^-^/H^+^ exchangers, were found in the ER membrane colocalized with SERCA. The ClC as well as SERCA mediated proton efflux must be compensated in some way in order to maintain the luminal pH neutralization. Early studies provided evidence for the existing proton influx towards the ER/SR lumen (14, 15) and it was estimated that around 10% of the countercurrent needed for compensating charge imbalance during Ca^2+^ release comes from proton movements (16, 17). However, this ER proton intake mechanism remains unclear (18). One possible contributor could be the K^+^/H^+^ exchangers (11) while Na^+^/H^+^ exchangers or Ca^2+^-leak channels are also potential candidates (18).

Recent studies on SERCA’s regulation have shed new light on the proton intake into the ER/SR membrane (19, 20). SERCA is prominently regulated by a transmembrane 52-residue-long protein, phospholamban (PLB), which binds and inhibits SERCA’s Ca^2+^ pumping activity (21). Structural and computational studies (22-24) have suggested the calcium regulation mechanism originates from the stabilization of a Ca^2+^-free E1 intermediate state of SERCA. Two protons were predicted to be in the binding-site and bind to E771 and E908 according to empirical pKa estimations. A transient-water-occupied pore was also identified by classical molecular dynamics (MD) simulations (19, 20, 24). This observation suggested the presence of a proton transport (PT) pathway for the release of a proton from residue E908 in the binding site to an intermediate luminal residue H944 and down to the luminal environment, However, classical MD simulations treat excess proton, water molecules, and protein in a non-reactive, fixed bonding topology manner with a fixed charge distribution, thus ignoring the well-known Grotthuss shuttling mechanism (25-28) of PT, the delocalization of the net positive excess protonic charge defect (29-31), and the altered hydration introduced by an explicit excess proton in the water structures (32-35). Alternative approaches, such as the hybrid quantum mechanical/molecular mechanical (QM/MM) method, can provide a reactive description of excess proton solvation and transport in protein channels due to the explicit treatment of the electronic structure. However, the QM/MM method is computationally expensive and allows for sampling on the tens to hundreds of picosecond timescales, thus limiting its ability to carry out the extensive free energy sampling required to fully understand PT processes in proteins (36). To overcome these sampling limitations, the multi-scale reactive molecular dynamics (MS-RMD) method (36-40) (and its predecessor the Multistate Empirical Valence Bond, or MS-EVB, method) have been developed, which are three orders of magnitude more computationally efficient compared to QM/MM. The MS-RMD models are developed from and calibrated against QM/MM data by utilizing a force matching algorithm in a “machine learning” type methodology. The MS-RMD approach can efficiently and accurately simulate explicit PT in proteins, as mediated by water molecules and amino acids while including Grotthuss proton shuttling, see, e.g., refs (34-36, 41-45).

In this work, MS-RMD simulations are employed along with free energy sampling (umbrella sampling) to test the classical MD-based hypothesis that PT occurs in SERCA from residue E908 into the lumen of the sarcoplasmic reticulum (19, 20). The free energy profile (potential of mean force, or PMF) was computed for the excess proton migration as well as its coupling to the fluctuations in water hydration of the transport pathway. Recent work in our group has revealed that a two-dimensional (2D) PMF is required to fully understand the PT mechanism and pathway in many proteins (32-35, 42, 44, 45). The two collective variables (CVs) defining the 2D PMF coordinates are the excess proton net positive charge defect location along the PT pathway and the water hydration (number of water molecules) occupying that pathway. It has been universally found so far that the excess proton motion and the water hydration structures are intrinsically coupled, i.e., the water hydration alone in the absence of an excess proton in the “water wire” is not sufficient for understanding the PT pathway and its mechanism, nor the rate of the proton translocation along that pathway. From either the 1D or 2D PMF, the rate of PT can be estimated from transition state theory (TST) (46, 47). By explicitly calculating the 2D PMF for the excess proton translocation and its hydration, the results presented in this work provide a quantitative description and molecular detail of one PT mechanism through SERCA, and our results support the hypothesis that this passive PT occurs on the microsecond timescale.

## MATERIALS AND METHODS

### Classical equilibration

Classical MD was continued from the equilibrated configuration of SERCA from previous studies (19, 20, 24). The system was embedded in a lipid bilayer of 368 POPC lipids and solvated with 49412 water molecules and 115 K^+^ and 93 Cl^-^ ions in an 11.68 Å × 11.68 Å × 15.4577 Å simulation box. Molecular interactions were described using the CHARMM36 force field (48, 49) and the TIP3P water model (50). The temperature was controlled by a modified velocity rescale thermostat (51) at 310 K and the pressure was controlled by the Berendsen barostat (52) at 1 atm. The system was integrated with a 2-fs timestep with all bonds involving hydrogen atoms constrained by the LINCS algorithm (53). The short-range interactions were smoothly switched off between 10 Å and 12 Å using the force-switching scheme and the long-range interactions were computed by the particle mesh Ewald method. The classical simulation was carried out with the GROMACS MD package (54, 55).

### MS-RMD method and model development

The MS-RMD methodology is described in more detail elsewhere (36-38, 40). Here we outline the essential aspects of MS-RMD. A reactive molecular system is described by a quantum-like Hamiltonian:

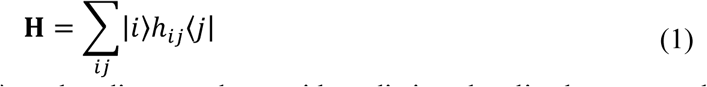

Each basis state |*i*⟩ corresponds to a given bonding topology with a distinct localized protonated species. The reactive process (change in bonding topology) and excess proton charge defect delocalization is described using a linear combination of basis states so that the ground state of the system |*ψ*⟩ is expanded in the basis set {|*i*⟩}:

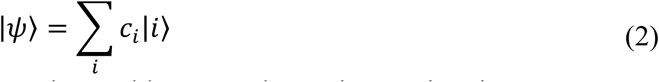

The coefficients *c*_*i*_ are solved from the eigen-value problem at each MD integration timestep:

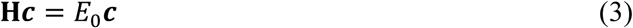

The diagonal terms *h*_*ii*_ in **H** can be described by a fixed bonding topology molecular mechanics (MM) potential using, in the present case, a modified version of the CHARMM36 force field (39). The classical force field is not capable of reactive simulations, so *h*_*ii*_ also includes an additional correction for shifting the diabatic surfaces referred to different zero points (38, 39). The off-diagonal matrix terms *h*_*ij*_, which provide the proton transfer mechanism between waters and waters with amino acid was chosen as the same functional form as previous studies, see ref (37, 56) for the amino acids and also the Supporting Material for details. The excess proton in the water molecules was described by the MS-EVB 3.2 model (40). The parameters in the Glu/His-water off-diagonal and the diabatic corrections for protonated Glu and His states were fit to QM/MM forces by minimizing the force residual 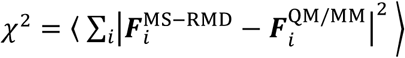. The fitted parameters were summarized in Table S1.

The configurations used in the training set were sampled by MS-RMD umbrella sampling biasing the distance between the excess proton center of excess charge (CEC, as defined below in Eq. 4) and the proton-acceptor atoms in Glu and His, namely the carboxylic oxygens of Glu and the imidazole nitrogen atoms of His. For E908, the umbrella windows range from 1.75 Å to 4.00 Å with a 0.25 Å separation. For H944, the umbrella windows range from 1.25 Å to 4.25 Å separated by a 0.25 Å spacing. The configurations were collected every 2 ps in each window, resulting in ∼1000 frames in total in both cases. Single-point QM/MM calculations were performed on the sampled configurations as reference forces. The MM part in QM/MM calculation was described by the standard CHARMM36 force field. The QM part was described by density functional theory at the BLYP/TZV2P level of theory. Besides the two titratable residues E908 and H944, sidechains of T799, V798, W794, V905, M909, S940, T763, V795, S767, S902, and N911 were all included as QM atoms. The electronic structure of three solvation shells of water around E908, H944, T799, T763, S767, and W794 were also treated explicitly. The QM/MM electrostatic coupling was computed by the Gaussian Expansion of the Electrostatic Potential (GEEP) method with periodic boundary conditions (57, 58). The broken alpha carbon and beta carbon bonds that crossed the QM/MM boundaries were capped with hydrogen atoms and the QM and MM forces were incorporated by the IMOMM scheme (59) with a scaling factor of 1.50.

### Umbrella sampling molecular dynamics

The MS-RMD umbrella sampling was conducted using a modified version (60) of the LAMMPS MD package (61) with PLUMED2 (62). The non-bonded interactions were truncated at 10 Å and the long-ranged interactions were computed by the particle-particle particle-mesh (PPPM) algorithm (63). The system was integrated using a 1-fs timestep by the Nose-Hoover chain thermostat (64) at 310 K under constant volume. The position of the excess proton net positive charge defect was defined by the center of excess charge (the important notation here and what follows given by “CEC”) (65):

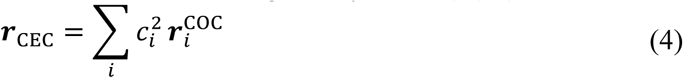

where *c*_*i*_’s are the ground state coefficients of the MS-RMD Hamiltonian and ***r***_COC_ is the center of charge of the species that holds the excess proton in basis state |*i*⟩. The CEC mathematically defines the location of the dynamical “electron hole”, which is the net positive charge defect caused by having an excess proton in the system (e.g., a hydrated excess proton defect that is Grotthuss shuttling through a chain or “wire” of water molecules and possibly also through protonatable amino acids).

In order to define the proton pathway, a short metadynamics (MTD) (66) run (∼3 ns) was conducted using the following collective variable:

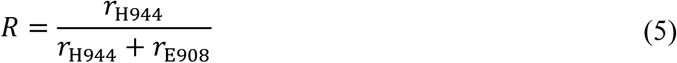

where *r*_H944_ indicates the distance between CEC and H944 and *r*_E908_ is the distance between CEC and E908. The Gaussian height was 0.25 kcal/mol with *σ* = 0.01 and placed every 1 ps. During the MTD, 4-5 transitions between E908 and H944 were observed and Hastie’s algorithm (67) was used to extract the principal curve through the point cloud formed by the CEC positions sampled. The projection of the CEC onto the curve (68) was then defined as the collective variable (CV) used in the production runs of umbrella sampling for the proton transport between E908 and H944. The proton release pathway from H944 to the lumen is relatively straightforward and thus the CV was defined as the distance between the excess proton CEC and H944, as projected on the averaged vector pointing from H944 nitrogen moiety towards the water in the pore between H944 and the lumen. The hydration CV in the 2D PMFs was taken as the water occupancy number (32) between E908 and H944 and between H944 and the lumen, respectively. Each umbrella sampling window was run for 150 ps to 4 ns, depending on convergence, resulting in a cumulative simulation time of 0.6 *μ*s.

### PT rate calculation

The PT rate calculation was based on transition state theory (46, 47):

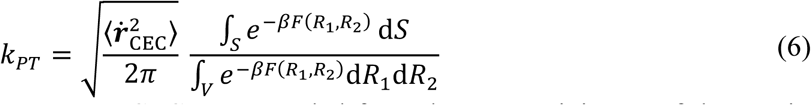

where the velocity of the excess proton CEC was sampled from the PMF minimum of the starting (reactant) state, the denominator is a two-dimensional integral over the reactant basin with respect to the two CVs (CEC position and water hydration occupancy, denoted here as *R*1 and *R*2), and the integral in the numerator was performed over the one-dimensional dividing surface at the transition state.

## RESULTS AND DISCUSSION

Previous microsecond-long classical MD simulations have shown that in the presence of bound PLB, SERCA populates a metal-ion-free E1 state (Fig. 1A) where transport site residues E771 and E908 are protonated (24). Based on these studies, a transient water pathway connecting residues E908 and H944 was seen in SERCA (Fig. 1B). This water pathway runs through transmembrane helices TM6, TM8, and TM9, and was stable for about 100 ps (19). Here, we used the microsecond-equilibrated structure reported in these studies as a starting structure to investigate the hydration environment of the pore from H944 to the lumen formed by transmembrane helices TM8, TM9, and TM10 (Fig. 1B). During the 200 ns of simulation, the pore between E908 and H944 showed only transient solvation, while the hydration of the pore between H944 and bulk was much more stable yet not always hydrated. Owing to these distinct hydration profiles of the two pores found in this study, we split the whole PT process into two steps (i) PT from E908 to H944, and (ii) from H944 down to the lumen.

**Figure 1.**
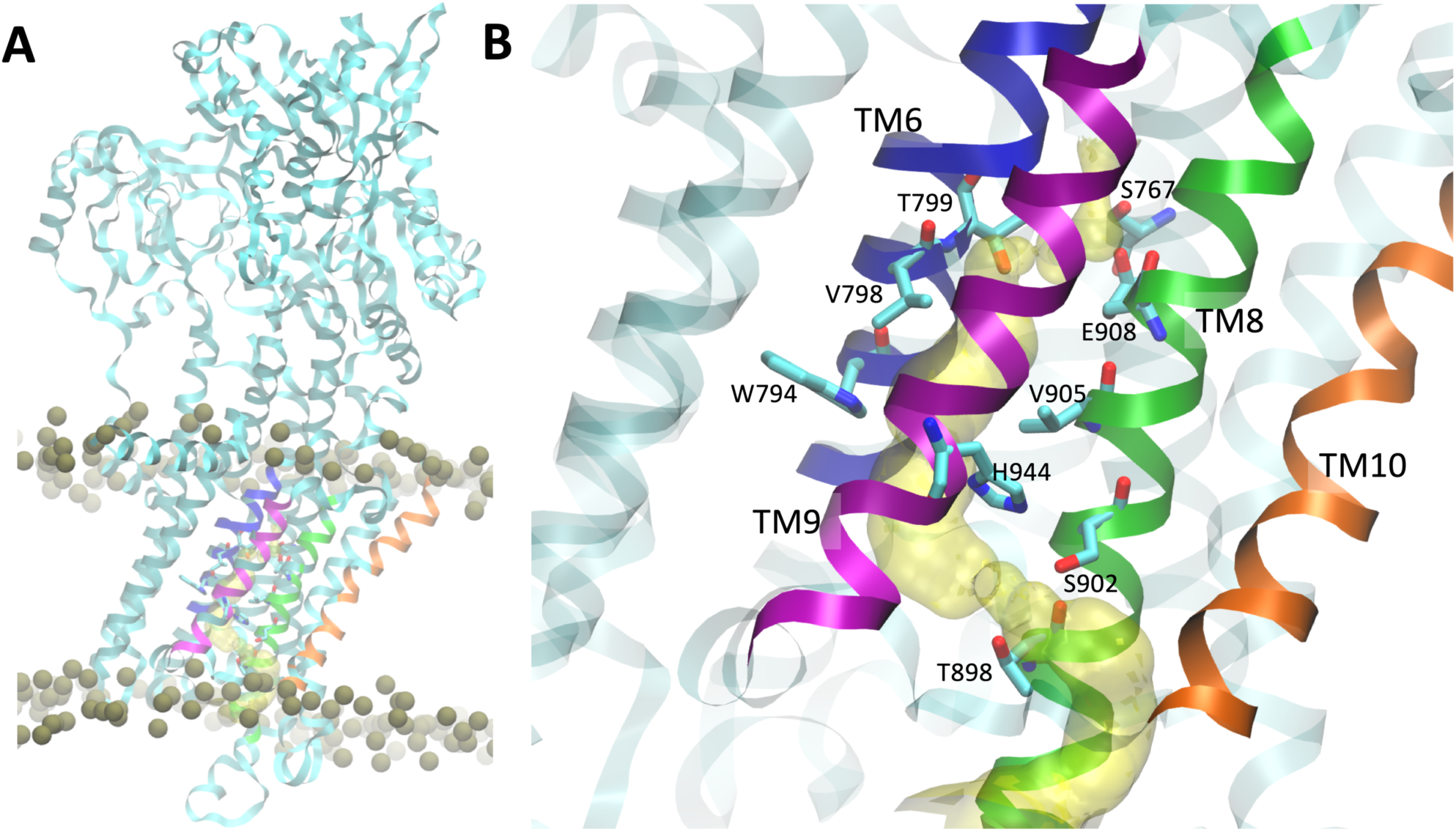
Location of the luminal water pore of SERCA. (A) Structure of SERCA bound to PLB and embedded in a lipid bilayer. The proteins are shown as ribbons, and the phosphate groups of the lipids are shown as gold spheres. For clarity, the TM helices are colored as follows: TM6, blue; TM8, green; TM9, purple; TM10, orange. (B) Luminal water pore of SERCA located between transmembrane helices TM6, TM8, TM9, and TM10; the pore is shown as a yellow surface.

### Proton transport from E908 to H944

The 2D PMF (Fig. 2A) for PT from E908 to H944 reveals a very clear coupled mechanism of the excess proton CEC movement with the pathway hydration for the overall PT mechanism. By following the minimum free energy path (black curve, the MFEP) in Fig. 2A, it is seen that the channel first becomes hydrated (movement in the vertical direction) to provide a hydrated electrostatic environment and adequate hydrogen bonds to help the breakage of the hydrogen bond between residues E908 and S767 (i.e., transition A→B, Fig. 3). The coordinated waters also allow charge defect delocalization of protonated E908, facilitating the proton dissociation from that residue. At the same time, the hydrated excess proton draws more water into the channel, inducing an increase in water occupancy between step B and the formation of the transition state (T) of the PT process (Fig. 3). The free energy barrier up to the transition state arises from two contributions: the deprotonation of E908 and unfavorable protonic charge localization induced by poor hydration of hydrophobic residues V798 and V905. This finding is clearly illustrated in Fig. 3T, showing that water molecules above or below the two hydrophobic gating residues are either hydrogen-bonded to other water molecules or to residues T799 and H944, contrary to the less solvated waters close to residues V798 and V905. This solvation imbalance results in a tendency for the proton to go back towards E908 or head to H944 and thus makes the transition state overlapping with the positions of V798 and V905. After overcoming the barrier, the system releases 5.9 kcal/mol of free energy from protonating H944 (state C). Then the channel becomes dehydrated (transition C→D), causing the free energy to decrease by 1.9 kcal/mol. The total reaction barrier for PT from E908 to H944 (A→T) is 8.7±0.3 kcal/mol, corresponding to a rate constant of 0.70±0.08 *μ*s^−1^, indicating that PT occurs through this pathway on the microsecond time scale, in agreement with the suggestion that passive proton transport can occur via this pore (20).

**Figure 2:**
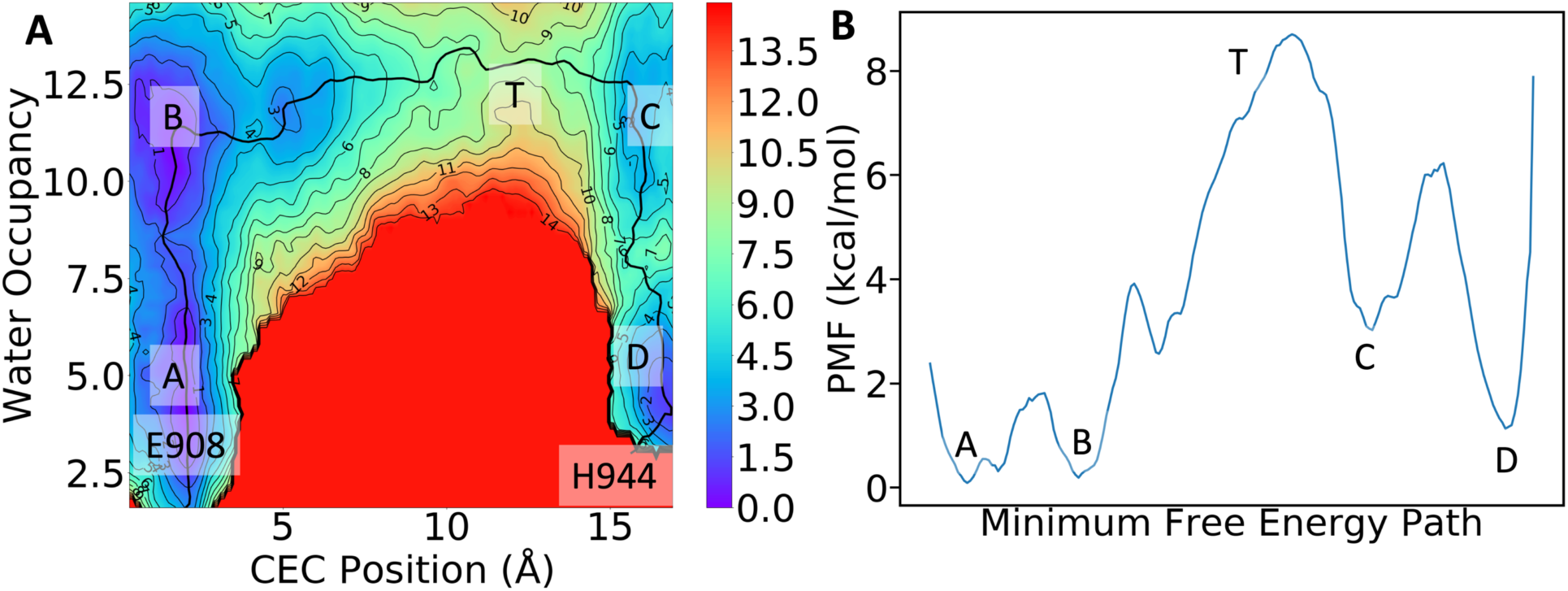
(A) Two-dimensional potential of mean force for proton transport from E908 to H944. (B) Free energy along the minimum free energy path (MFEP).

**Figure 3:**
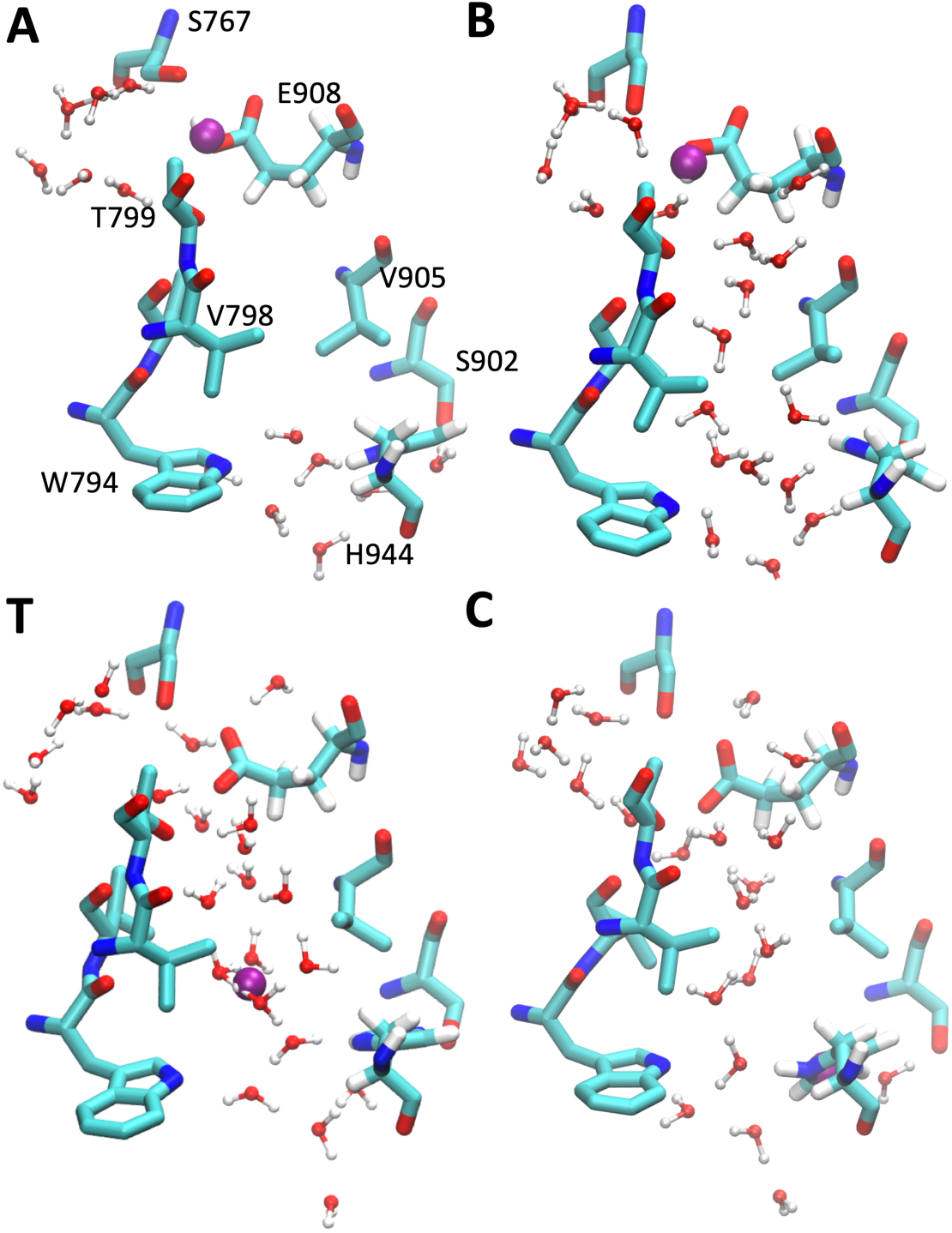
Representative configurations along the minimum free energy path, with labels showing the positions on the 2D PMF (Fig. 2A). The position of the excess proton defect CEC is rendered as a purple sphere. The hydrogen atoms of S767, T799, V798, V905, S902, and W794 are not shown for clarity. (A) The protonated E908 forms a hydrogen bond with S767. (B) The channel becomes hydrated and the E908-S767 hydrogen bond breaks. (T) The transition state of the PT reaction where the excess proton is solvated in the water close to the hydrophobic V798 and V905 residues. (C) The excess proton shuttles to H944 via water wires.

### Proton transport from H944 to the lumen

We next focused on the second PT step corresponding to proton permeation from H944 to the luminal bulk via the pore formed by transmembrane helices TM8, TM9, and TM10. We found in our classical MD equilibration that the pore is better hydrated due to the hydrophilic environment created by residues S902 and T898. Due to this hydrophilic pore nature, the 2D PMF of excess proton CEC motion and water occupancy for this PT step (Fig. 4A) follows a simpler mechanism compared to the PT from E908 to H944 reported in the prior section. Specifically, the PMF features a free-energy well corresponding to a protonated H944 and solvation water number ranging from 5 to 7. The transition state is located 3.6 Å away from H944, which coincides with the position of the bottleneck of the pore formed by residues S902 and T898. As shown in Fig. 4C, around the transition state the two gating residues, S902 and T898, replace two waters in the first and second solvation shell of the proton-water motif. As Ser and Thr are less basic than water, they destabilize the excess proton solvation and increase the free energy in this region. Compared to the gating residues V798 and V905, the S902 and T898 gate results in a much lower barrier of 3.9±0.4 kcal/mol. From the transition state theory, we obtained a rate constant of 3.1 ±1.5 ns^-1^. Compared to the microsecond timescale of the previous PT step, this nanosecond timescale indicates the proton can easily exit the protein after it reaches H944 from E908. Based on these findings, we propose that the E908→H944 PT step in the previous section serves as the rate-limiting step of the entire PT process.

**Figure 4:**
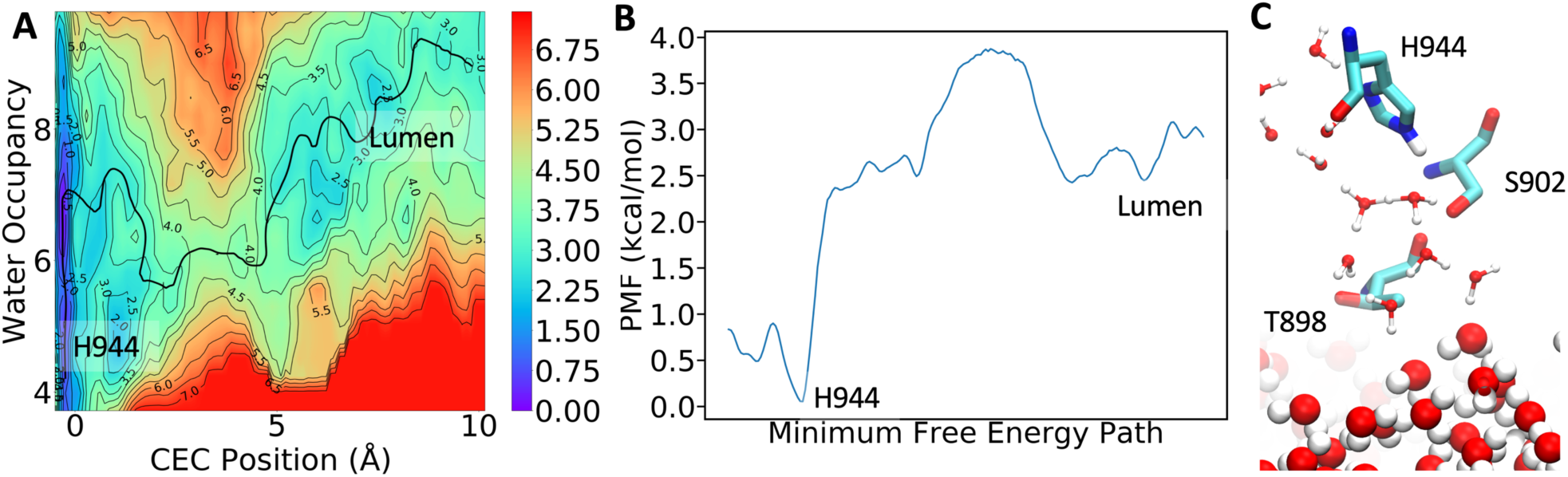
(A) 2D PMF for PT from H944 to the lumen. (B) Free energy along the MFEP. (C) Representative configuration at the transition state.

## CONCLUSIONS

Extensive multiscale simulations have been reported in this paper to study the PT process through SERCA. To the best of our knowledge, this is the first computational study of this large biomolecular system that has included the explicit physical process of proton transport via Grotthuss shuttling. The efficiency of the MS-RMD simulation approach has enabled the simulation of the proton translocation along a 25 Å-long pathway at atomistic-level detail for nearly a microsecond of total simulation time, which is well beyond the achievable timescale of QM/MM simulation. The simulations included the calculation of 2D PMFs for the complex PT process, revealing the coupled role of hydration with the excess proton translocation in the pathway. The calculated rate constant along the minimum free energy path of the 2D PMF reveals a microsecond timescale for proton transport from the Ca^2+^-binding site to the lumen. This result thus highlights the crucial role of a pore that was discovered in a Ca^2+^-free E1 state of SERCA for deprotonating the binding site and reactivating SERCA into the Ca^2+^-affinitive E2 state. More importantly, the pore was shown to be a feasible passive proton transport pathway mediated by SERCA that may explain the proton flux towards the SR/ER lumen observed in experiments. The PT process involves the breakage of the hydrogen bond between S767 and E908 and proton permeation through the V798-V905 gate. It is thus proposed that the residues S767, V798, and V905 – which are conserved in the SERCA family – are possible targets for future experimental mutagenesis studies.

## AUTHOR CONTRIBUTIONS

G.A.V. and L.M.E.-F. designed research; C.L. and Z.Y. performed research; C.L., Z.Y., G.A.V., and L.M.E.-F. analyzed data; C.L., Z.Y., G.A.V., and L.M.E.-F. wrote the paper.

## ACKNOWLEDGMENTS

The personnel in this research were supported by the National Institute of General Medical Sciences (NIGMS) of the National Institutes of Health (NIH) through grants R01 GM053148 (to C.L., Z.Y. and G.A.V) and R01 GM120142 (to L.M.E.-F.). The computational resources in this research were provided by the Extreme Science and Engineering Discovery Environment (XSEDE), which is supported by the National Science Foundation Grant Number ACI-1053575, and the University of Chicago Research Computing Center (RCC). We thank Professor Jessica Swanson for many enlightening discussions and her valuable input, as well as Paul Calio for reading the manuscript and providing helpful comments.

