## Supplemental File for "Multiscale Simulation Reveals Passive Proton Transport Through SERCA on the Microsecond Timescale"

### Supporting Information for
### Multiscale Simulation Reveals Passive Proton Transport Through SERCA on the Microsecond Timescale

Chenghan Li,[1] Zhi Yue,[1] L. Michel Espinoza-Fonseca,[2] and Gregory A. Voth[1]

[1]Department of Chemistry, Chicago Center for Theoretical Chemistry, James Franck Institute, and Institute for Biophysical Dynamics, The University of Chicago, Chicago, Illinois 60637, United States
[2]Center for Arrhythmia Research, Department of Internal Medicine, Division of Cardiovascular Medicine, University of Michigan, Ann Arbor, Michigan 48109, United States

**The functional form of the MS-RMD models**

The off-diagonal interaction used in this work between Glu and water was

$$h_{ij} = V_{ij}^{\mathrm{const}} \cdot \exp(-\gamma \boldsymbol{q}^2) \cdot \exp\big(-\alpha(|\boldsymbol{r}_O - \boldsymbol{r}_{O'}| - R_{OO}^0)\big)$$

where $\boldsymbol{q} = (\boldsymbol{r}_O + \boldsymbol{r}_{O'})/2 - \boldsymbol{r}_{H^*}$ is the asymmetric stretch coordinate ($\boldsymbol{r}_O$ and $\boldsymbol{r}_{O'}$ are the positions of the carboxylic oxygen of Glu and the water oxygen, and $\boldsymbol{r}_{H^*}$ is the shared proton of the two oxygens). The diabatic correction added on the protonated Glu state was a constant, $V_{ii}^{\mathrm{const}}$, while the correction for deprotonated Glu state was Eqs. 7-9 in ref (1).

The off-diagonal between His and water was Eqs. 4-8 in ref (2). The diabatic correction for protonated His state and deprotonated state are both constants following the ref (2), denoted as $V_{ii}^{\mathrm{HSP}}$, $V_{ii}^{\mathrm{HSD}}$ and $V_{ii}^{\mathrm{HSE}}$ respectively in Table S1.

**TABLE S1. The MS-RMD model parameters.**

A)

| E908 parameters | |
|---|---|
| $V_{ij}^{\mathrm{const}}$ | -27.754 |
| $\gamma$ | 1.3056 |
| $\alpha$ | 0.0392 |
| $R_{OO}^0$ | 3.5999 |
| $V_{ii}^{\mathrm{const}}$ | -145.84 |

B)

H944 off-diagonal parameters

| | N-$\epsilon$ | N-$\delta$ |
|---|---|---|
| $V_{ij}^{\mathrm{const}}$ | -31.177 | -12.854 |
| $r_{\mathrm{sc}}^0$ | 1.3056 | 1.2057 |
| $\lambda$ | 0.59978 | 0.31334 |

| | | |
|---|---|---|
| $R_{DA}^0$ | 2.6544 | 2.4549 |
| $C$ | 0.57498 | 0.68372 |
| $\alpha$ | 1.2677 | 0.32141 |
| $a_{DA}$ | 2.7994 | 2.3469 |
| $\beta$ | 0.25061 | 0.42373 |
| $b_{DA}$ | 1.9964 | 2.0944 |
| $\epsilon$ | 1.2459 | 12.972 |
| $c_{DA}$ | 3.1494 | 1.0983 |
| $\gamma$ | 1.8713 | 4.0248 |

C)

| H944 diabatic corrections | |
|---|---|
| $V_{ii}^{\mathrm{HSP}}$ | -102.41 |
| $V_{ii}^{\mathrm{HSD}}$ | -12.700 |
| $V_{ii}^{\mathrm{HSE}}$ | 0 |
